# Genome-wide patterns of *de novo* tandem repeat mutations and their contribution to autism spectrum disorders

**DOI:** 10.1101/2020.03.04.974170

**Authors:** Ileena Mitra, Bonnie Huang, Nima Mousavi, Nichole Ma, Michael Lamkin, Richard Yanicky, Sharona Shleizer-Burko, Kirk E. Lohmueller, Melissa Gymrek

## Abstract

Autism Spectrum Disorder (ASD) is an early onset developmental disorder characterized by deficits in communication and social interaction and restrictive or repetitive behaviors^1,2^. Family studies demonstrate that ASD has a significant genetic basis^3^ with contributions both from inherited and *de novo* variants. While the majority of variance in liability to ASD is estimated to arise from common genetic variation^4^, it has been estimated that *de novo* mutations may contribute to 30% of all simplex cases, in which only a single child is affected per family^5^. Tandem repeats (TRs), consisting of approximately 1-20bp motifs repeated in tandem, comprise one of the largest sources of *de novo* mutations in humans^6^. Yet, largely due to technical challenges they present, *de novo* TR mutations have not yet been characterized on a genome-wide scale, and their contribution to ASD remains unexplored. Here, we develop novel bioinformatics tools for identifying and prioritizing *de novo* TR mutations from whole genome sequencing (WGS) data and use these to perform a genome-wide characterization of *de novo* TR mutations in ASD-affected probands and unaffected siblings. Compared to recent work on TRs in ASD^7^, we explicitly infer mutation events and their precise changes in repeat copy number, and primarily focus on more prevalent stepwise copy number changes rather than large or complex expansions. Our results demonstrate a significant genome-wide excess of TR mutations in ASD probands. TR mutations in probands tend to be larger, enriched in fetal brain regulatory regions, and predicted to be more evolutionarily deleterious compared to mutations observed in unaffected siblings. Overall, our results highlight the importance of considering repeat variants in future studies of *de novo* mutations.

## Identifying de novo TR mutations

We developed a novel method, MonSTR, for identifying *de novo* TR mutations in parent-offspring trios from whole-genome sequencing (WGS) data (**Methods; Supplementary Methods**). MonSTR takes genotype likelihoods reported by a TR genotyper as input and estimates the posterior probability of a mutation resulting in a repeat copy number change at each TR in each child.

We performed a genome-wide analysis of *de novo* TR mutations (Fig. 1a) using WGS available for 1,637 quad simplex families sequenced to 35× coverage as part of the Simons Simplex Collection^8^ (SSC) (**Supplementary Table 1**), which have been ascertained to enrich for probands likely to harbor previously uncharacterized pathogenic *de novo* mutations^9^. We used GangSTR^10^ to estimate diploid repeat lengths in each sample at 1,189,198 TRs with repeat unit lengths 1-20bp and median total lengths 12bp in hg38. TR genotype results were used as input to MonSTR to identify mutations in each child. After filtering (**Methods**), 1,593 families remained for analysis. Our pipeline identified a total of 175,291 high-confidence TR mutations across 94,616 distinct loci (average 53.9 autosomal mutations; Fig. 1b) corresponding to an average mutation rate of 5.6×10^-5^ mutations per locus per generation.

**Figure 1:**
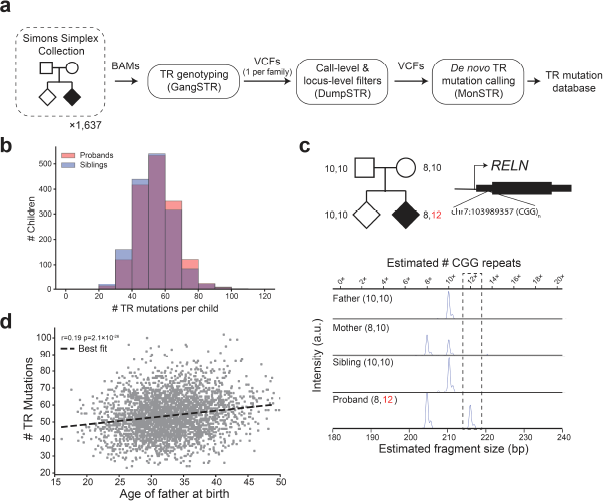
Identifying *de novo* TR mutations in the SSC cohort. **a. Overall study design**. We analyzed *de novo* TR mutations from WGS data of 1,637 quad families consisting of father, mother, non-ASD child (sibling), and ASD affected child (proband) from the Simons Simplex Collection. After filtering (**Methods**), 1,593 families remained for analysis. **b. Distribution of the number of *de novo* TR mutations for each child**. The histogram displays the distribution of autosomal *de novo* TR mutation counts for non-ASD siblings (blue) and probands (red). **c. Example TR mutation validated by capillary electrophoresis**. A mutation resulting in two additional copies of CGG in the 5’UTR of the gene *RELN* (chr7:103989357; hg38) was identified based on WGS analysis (top). Alleles with 11 or more copies of CGG at this TR were previously implicated in ASD^57^ and have been shown to reduce *RELN* expression^58^. The region was amplified by PCR and the mutation was confirmed by capillary electrophoresis. Estimated fragment sizes for each sample (bottom x-axis) and the corresponding repeat numbers (top x-axis) are annotated. The fragment length corresponding to the *de novo* allele (12×) is denoted by the dashed gray box. **d. Correlation of mutation rate with paternal age**. The scatter plot shows the father’s age at birth (x-axis) vs. the number of autosomal *de novo* TR mutations identified per child (y-axis). Each gray point represents one child (n=3,186). The dashed black line gives the best fit line.

We tested our framework on simulated WGS data, which demonstrated high sensitivity to detect TR mutations resulting in changes of up to 10 repeat copies and low false positive rate (<1%) compared to a naïve method in most settings (**Methods**, Extended Data Fig. 1). To directly assess the quality of mutation calls in SSC, we performed fragment analysis using capillary electrophoresis (CE) on a subset of 49 TR mutations across 5 SSC quad families (example in Fig. 1c). In total, we tested 196 individual genotypes and 98 transmission events (Supplementary Tables 2-3). Tested mutations show a validation rate of 90% (44/49), an improvement over validation rates previously reported for *de novo* indel calling^11^. To further validate MonSTR mutation calls, we compared our results to known TR mutation trends (Extended Data Fig. 2). Similar to previous studies^12–14^, estimated mutation rates are highest for TRs with shorter repeat units (Extended Data Fig. 2a) and are positively related to total length (bp) of the of the reference TR (Extended Data Fig. 2b). Following *de novo* single nucleotide variant (SNV) studies^9,15^, autosomal TR mutation rates are correlated with paternal age (Pearson *r*=0.19; two-sided p=2.1×10^-26^; n=3,186; Fig. 1d). At TR mutations (excluding homopolymers) for which the parent of origin could be inferred (**Methods**), 74% were phased to the father, which is similar to previous reports for *de novo* SNVs^16,17^. Finally, genome-wide observed mutation counts in SSC are significantly correlated with rates estimated by our MUTEA^12^ method on an orthogonal set of unrelated individuals (Pearson r=0.26; p<10^-200^; n=548,724; Extended Data Fig. 2d). Taken together, these results suggest our pipeline can robustly identify genome-wide *de novo* TR mutations.

## Genome-wide patterns of TR mutations

We first characterized genome-wide properties of autosomal TR mutations. The majority of mutations observed result from expansions or contractions by a single repeat unit, with a smaller number of larger mutations (Fig. 2a). Mutation step size distributions vary noticeably by repeat unit length (**Supplementary Table 4**, Extended Data Fig. 3a), which has been previously shown^14,18–20^. Overall, mutations show a bias toward expansions (71%) vs. contractions (29%). When excluding error-prone homopolymer TRs, only 56% of mutations are expansions, still significantly more than the 50% expected by chance (binomial two-sided p=4.8×10^-249^; n=71,822).

**Figure 2:**
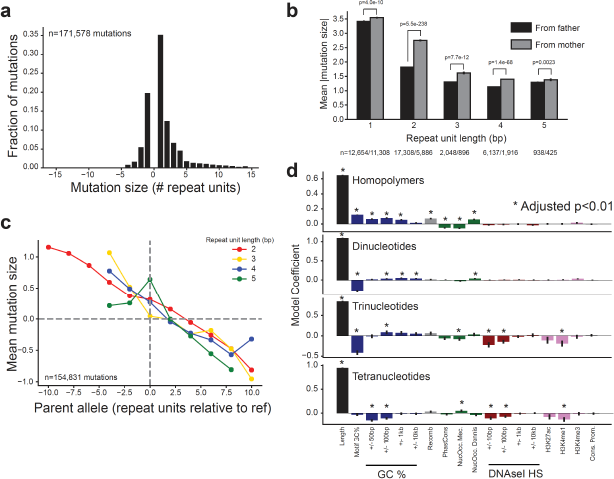
Genome-wide patterns of TR mutations. **a. Mutation size distribution**. The histogram shows the distribution of mutation sizes in terms of the number of repeat units, where mutations >0 represent expansions and <0 represent contractions. **b. Mean absolute mutation size by parental origin**. Bars show the mean absolute mutation size for mutations phased to the paternal germline (black bars) and the maternal germline (gray bars). The x-axis denotes the length of the repeat unit in bp. One-sided p-values were computed using a Mann-Whitney test. **c. Directionality bias in mutation size**. The x-axis gives the size of the parent allele relative to the reference genome (hg38). The y-axis gives the mean mutation size across all mutations with given parent allele length. A separate line is shown for each repeat unit length (red=dinucleotides; gold=trinucleotides; blue=tetranucleotides; green=pentanucleotides). To reduce potential influence of heterozygote dropout errors (**Methods**), only mutations with step size of up to ±5 units are included. **d. Determinants of TR mutation rates**. The Poisson regression coefficient is shown for each feature in models trained separately for each repeat unit length. Features marked with an asterisk denote significant effects (p<0.01 after Bonferroni correction for the number of features tested across all four models).

We further examined mutation sizes separately for the subset of mutations phased to either the maternal vs. paternal germline. The bias toward expansions vs. contractions (excluding homopolymers) is significant for maternal mutations (57% expansions; binomial two-sided p=3.7×10^-39^; n=9,190) but not for paternal mutations (50% expansions; p=0.71; n=26,550) (Extended Data Fig. 3b-c), suggesting the overall expansion bias observed is primarily driven by maternally derived mutations. Further, maternal mutations result in significantly larger changes in repeat unit copy number (Mann-Whitney one-sided p<10^-200^; maternal median size=2 units, paternal median size=1 unit). This trend is recapitulated across all repeat unit lengths (Fig. 2b), with the strongest effect at dinucleotide TRs.

Previous studies assessing TR patterns reported a directionality bias in mutations, with longer alleles more likely to experience contractions and shorter alleles more likely to experience expansions^12,14,21^. We analyzed mutation sizes at TRs with unit lengths from 2-5bp and observe a similar bias (Fig. 2c). We find that the directionality bias is notably stronger for mutations originating from parents heterozygous for two different allele lengths (Extended Data Fig. 3d-e), whereas little bias is observed for mutations from homozygous parents. This suggests the observed trend could be driven in part by interaction between parent alleles, which has been previously hypothesized^21^.

Finally, we investigated relationships between TR mutation rates and genomic or epigenomic features (**Methods**). We fit a separate Poisson regression for each repeat unit length relating observed mutation counts to each feature. As expected, reference TR length is the strongest predictor of mutation rates across all TRs (Fig. 2d). Several features show similar patterns across all TRs, including the presence of active chromatin marks (negative effect) and recombination rate (positive effect, which has been previously suggested^22^). Other features, such as GC content, show distinct patterns across different TR classes. These results suggest TR variation is driven by a variety of mutational mechanisms that may be unique to each TR unit class.

## TR mutation burden in ASD

The total number of *de novo* autosomal TR mutations observed genome-wide is significantly higher in probands (mean=54.65 mutations) vs. non-ASD siblings (mean=53.05 mutations) (Fig. 3a, paired t-test two-sided p=9.4×10^-7^; n=1,593; Relative risk [RR] = 1.03). This trend remains after adjusting mutation counts for paternal age (p=1.08×10^-5^; **Methods**), excluding homopolymers (p=0.0071 after paternal age adjustment), and is consistently observed across each SSC phase (**Supplementary Table 5**). Autosomal mutations in probands result in significantly larger repeat copy number changes (Mann Whitney one-sided p=0.017; Fig. 3b). We analyzed chromosome X mutations separately and observed a moderate excess in male probands vs. male non-ASD siblings (Mann-Whitney two-sided p=0.01) but not in females (p=0.69).

**Figure 3:**
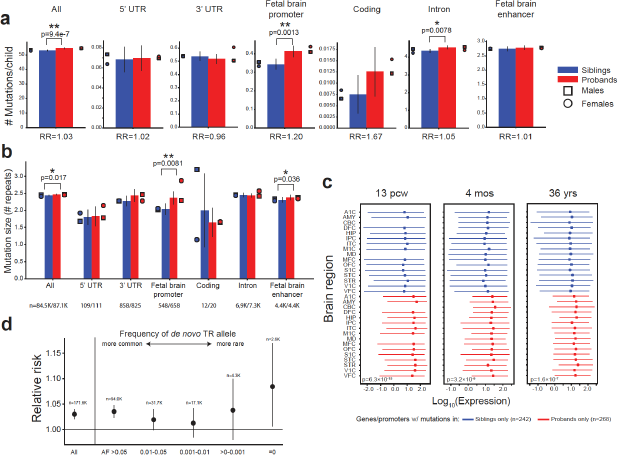
TR mutation burden in ASD. **a. Mean mutation counts in probands vs. non-ASD siblings by gene annotation**. Blue bars denote non-ASD siblings, red bars denote probands. Circles and squares show counts for females and males, respectively. **b. Mutation sizes in probands vs. non-ASD siblings**. Bars denote mean mutation sizes in terms of the absolute number of repeat units. In **a-b**, error bars give 95% confidence intervals. Single and double asterisks denote significant increases (p<0.05) before and after Bonferroni correction, respectively. **c. Distribution of brain expression of genes with *de novo* TR mutations**. Red lines show the distribution of gene expression for genes with only proband mutations. Blue lines show distributions for genes with only mutations in non-ASD siblings. Dots give median expression and lines extend from the 25th to 75th percentiles (A1C=primary auditory cortex; AMY=amygdala; CBC=cerebellar cortex; DFC=dorsolateral prefrontal cortex; HIP=hippocampus; IPC=posterior inferior parietal cortex; ITC=inferolateral temporal cortex; M1C=primary motor cortex; MD=mediodorsal nucleus of thalamus; MFC=anterior cingulate cortex; OFC=orbital frontal cortex; S1C=primary somatosensory cortex; STC=posterior superior temporal cortex; STR=striatum; V1C=primary visual cortex; VFC=ventrolateral prefrontal cortex). P-values are based on a meta-analysis across brain regions (**Methods**). **d. Mutation burden by allele frequency (AF)**. The x-axis stratifies mutations based on non-overlapping bins of the frequency of the *de novo* allele in SSC parents. “All” includes all mutations. For other bins, only TRs for which precise copy numbers could be inferred in at least 80% of SSC parents are included. The y-axis gives relative risk (RR) in probands vs. non-ASD siblings.

Our study is underpowered to detect specific TR loci enriched for mutations in probands vs. siblings at genome-wide significance (Extended Data Fig. 4). Instead, we evaluated whether TRs within particular categories show an excess of mutations in probands vs. non-ASD siblings (Fig. 3a). Mutations in coding regions have the highest magnitude of excess in probands vs. non-ASD siblings, but the excess is not statistically significant (RR=1.67; paired t-test two-sided p=0.16) likely due to the small number of autosomal coding mutations (n=32, **Supplementary Table 6**). Similar to previous studies of non-coding point mutations^9,23^, we observe significant enrichment for *de novo* TR mutations falling within annotated fetal brain promoters (Fig. 3a; RR=1.20; paired t-test two-sided p=0.0013; significant after multiple hypothesis correction) and within 50kb of SNPs previously associated with schizophrenia and educational attainment (Extended Data Fig. 5; **Supplementary Note**). We examined expression of genes with coding or promoter mutations and found that genes with TR mutations only observed in ASD probands show significantly higher prenatal expression compared to genes with mutations found in non-ASD siblings (p=6.3x-10^-15^ at 13 post-conceptional weeks [pcw]; **Methods**; Fig. 3c; Extended Data Fig. 6a). Further, proband mutations are predicted to more significantly alter expression of nearby genes in the brain compared to control mutations (**Supplementary Note**; Extended Data Fig. 6b).

The observed genome-wide excess of TR mutations in probands is modest (RR=1.03), suggesting that only a subset are pathogenic. Indeed, the majority (84%) of TR mutations result in alleles that are already common (allele frequency [AF] ≥1%) in unaffected parents within the SSC cohort and thus are likely to be benign. When we stratify our mutation burden analysis by the frequency of the mutant allele (Fig. 3d), we find that the mutation excess in probands increases for mutations resulting in rarer alleles, with the strongest effect at alleles unobserved (AF=0) in SSC parents (RR=1.12; paired t-test two-sided p=0.021). This pattern remains after excluding error-prone homopolymer TRs (Extended Data Fig. 7). We identified specific TR mutations in coding or promoter regions resulting in alleles unobserved in unaffected parents (n=12 in probands, 7 in siblings). This set includes multiple genes previously implicated in ASD or related neurodevelopmental disorders^24–29^ in addition to novel candidate genes (Extended Data Fig. 8; **Supplementary Note**).

## Prioritizing pathogenic TR mutations

We sought to further prioritize TR mutations based on their predicted deleterious effects. Metrics commonly used to annotate SNV mutations^30–32^ are not applicable to TRs, which tend to be multi-allelic and result in either non-coding mutations or in-frame indels. To overcome this challenge, we developed a novel population genetics framework, Selection Inference at Short TRs (SISTR) to measure negative selection against individual TR alleles. SISTR fits an evolutionary model of TR variation that includes mutation, genetic drift, and negative natural selection to empirical allele frequency data (per-locus frequencies of each allele length) to infer the posterior distribution of selection coefficients (*s*) at individual TRs (Extended Data Fig. 9). Parameter *s* can be interpreted as the decrease in reproductive fitness impact for each repeat unit copy number away from the population modal allele at a given TR. Testing our method on simulated datasets capturing a range of mutation and selection models, SISTR accurately recovers simulated values down to *s*=10^-4^, corresponding to strong or moderate selection, for most settings (Fig. 4a; Extended Data Fig. 10a-b). Notably, SISTR currently only handles TRs with repeat unit lengths 2-4bp. Full descriptions of the mutation and selection model and SISTR inference method are given in the **Supplementary Methods**.

**Figure 4:**
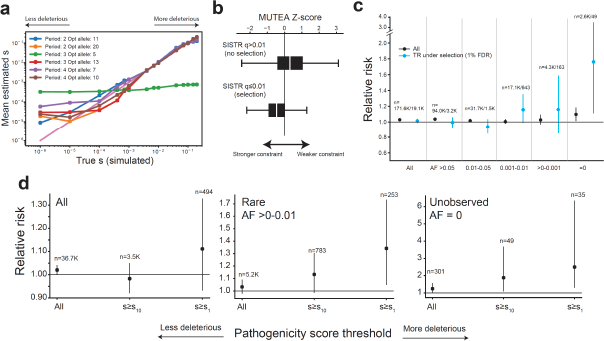
Prioritizing TR mutations by fitness effects. **a. Comparison of true vs. inferred per-locus selection coefficients**. The x-axis shows the true simulated value of *s*, and the y-axis shows the mean *s* value inferred by SISTR across 200 simulation replicates. Each color denotes a separate mutation model based on the mutation rate model for that period (repeat unit length; **Supplementary Methods**) and the length (in repeat units) of the optimal allele at that locus. **b. Comparison of SISTR and MUTEA**. Boxes show the distribution of MUTEA constraint scores for STRs inferred by SISTR to have non-significant (top) or significant (bottom) selection coefficients (FDR<1%). Negative MUTEA z-scores indicate stronger mutational constraint. White middle lines give medians and boxes span from the 25th percentile (Q1) to the 75th percentile (Q3). Whiskers extend to Q1-1.5*IQR (bottom) and Q3+1.5*IQR (top), where IQR gives the interquartile range (Q3-Q1). **c. Mutation burden at TR loci under negative selection**. The x-axis stratifies mutations based on the same allele frequency categories as in **a**. Blue dots give RR considering only TR loci inferred to be under the strongest negative selection (FDR<1%; **Methods**). **d. Per-allele selection coefficients further stratify mutation burden within allele frequency bins**. For mutations resulting in common (All TRs; left), rare (AF >0-0.01; middle) and unobserved (AF=0; right) TR alleles, we computed an allele-specific pathogenicity score based on inferred per-locus selection models (**Methods**). Larger *s* values denote a mutation resulting in an allele predicted to be more deleterious. *s_10_* and *s_1_* correspond to the top 10% and top 1% of pathogenicity scores, respectively. Error bars give 95% confidence intervals.

We applied SISTR to estimate selection coefficients at genome-wide TRs based on allele frequencies observed in unaffected SSC parents (**Supplementary Data 1**). After filtering (**Methods**), 82,223 STRs remained for analysis. We found that the overall distribution of selection coefficients is robust to input choices including demographic model and prior distribution on *s* (Extended Data Fig. 10c). We compared SISTR results to per-locus STR constraint scores we previously reported using MUTEA^12^. We found that STRs with significant selection coefficients have significantly stronger MUTEA constraint scores (Mann-Whitney one-sided p<10^-200^; Fig. 4b). We further compared SISTR results to gene-level constraint scores based on SNPs^31,33,34^. Protein-coding STRs under strongest negative selection tend to be in genes less tolerant of missense mutations (Mann Whitney one-sided p=0.00098; Extended Data Fig. 10d), or loss of function SNP mutations (Mann Whitney one-sided p=0.00219; Extended Data Fig. 10e), compared to coding STRs not inferred to be under negative selection (*s*=0).

We next tested for an enrichment of evolutionarily deleterious TRs in probands compared to non-ASD siblings. When restricting to TR loci predicted to be under selection (*s*>0 with false discovery rate [FDR] <1%), we find an increased mutational burden in probands (Fig. 4c), which is most notable for mutations resulting in rare *de novo* alleles. Stratifying mutations based on allele-specific selection coefficients results in a further increased mutational burden (Fig. 4d). *De novo* TR mutations with rare or unobserved allele frequencies and estimated to be the most deleterious (top 1% of *s* scores) show the strongest relative risk (RR=1.34 and RR=2.50 for rare [AF<0.01] or unobserved low fitness alleles, compared to RR=1.03 genome-wide). We identified 35 mutations, of which 25 are in probands, resulting in previously unobserved alleles predicted to be strongly deleterious (top 1% of *s* scores). Of these, multiple proband mutations are in genes with point mutations previously implicated in ASD (*e.g. PDCD1, KCNB1, AGO1, CACNA2D3, FOXP1, RFX3, MED13L*) or related phenotypes, whereas only two rare mutations are found in siblings to be related to ASD genes (**Supplementary Table 7**). Overall, these results suggest that the subset of TR mutations resulting in rare alleles under strongest selection are most pathogenic for ASD risk.

## Discussion

We present a novel framework for the identification and prioritization of *de novo* TR mutations. We analyze nearly 1 million TRs across more than 3,000 transmissions each and reveal on average 54 autosomal TR mutations per individual. The true number of mutations is likely underestimated due to the stringent filtering applied to candidate mutations. Overall, our results identify novel patterns of TR mutation (**Supplementary Discussion**) and suggest that the burden of *de novo* TR mutations is similar in magnitude to the total number of *de novo* point mutations per child^18,45^.

We find a significant genome-wide excess of *de novo* TR mutations in probands compared to non-ASD siblings. Based on this excess, we estimate that these mutations contribute to approximately 1.6% of simplex idiopathic ASD probands. A recent study analyzing an orthogonal set of variants estimated that larger complex TR expansions contribute to 2.6% of simplex cases. Taken together, these results suggest TRs may account for around 4% of simplex ASD cases, comparable in magnitude to non-coding point mutations^7^.

Importantly, only a subset of *de novo* TR mutations is likely to contribute to ASD risk or have deleterious effects. We find that mutations resulting in *de novo* alleles that are very rare (AF<0.001) or estimated to be under strong negative selection show the strongest signal of excess mutations in probands. The relative risk observed for these most severe mutations (RR=2.50), which are nearly all non-coding, is similar in magnitude to previously reported relative risks for protein-truncating variants^6^. On the other hand, we estimate the overall contribution to simplex ASD to be highest for mutations resulting in common alleles (of the 1.6% estimated above, 1.1% is attributed to mutations with AF>0.05). The impact of these mutations is not obvious and is the subject of future study.

Our study faced several limitations: (***i***) Identification of TR mutations remains challenging and requires stringent filtering to achieve high validation rates. (***ii***) Our results exclude important TR mutation classes, such as sequence interruptions^51^, somatic variation^52^, and complex repeat expansions which have been recently studied elsewhere^11^. (***iii***) We do not currently have power to implicate specific TRs at genome-wide significance (Extended Data Fig. 4). Future methods improvements and increasing sample sizes are likely to pinpoint specific TR mutations most relevant to ASD. The framework developed in our study will serve as a valuable resource for further characterizing TR mutations and their role in ASD and other diseases.

## Methods

### Dataset and preprocessing

The Simons Simplex Collection (SSC) dataset used in this study consists of 1,637 quad families (**Supplementary Table 1**). Informed consents were obtained for each participant by the respective studies in accordance with their local IRBs. Our study used only de-identified data, and thus was exempt from institutional review board (IRB) review by the University of California San Diego IRB (Project # 170840). Access to SSC data was approved for this project under SFARI Base project ID 2405.2. CRAM files containing WGS reads aligned to the hg38 reference genome and phenotype information for phases 1-3 were obtained from SFARI base (https://base.sfari.org/).

### Genome-wide TR genotyping

CRAM files were processed on Amazon Web Services (AWS) using the AWS Batch service. Genotyping of autosomal TRs was performed with GangSTR^10^ v2.4.2 using the reference TR file hg38_ver16.bed.gz available on the GangSTR website (https://github.com/gymreklab/GangSTR) and with the option --include-ggl to enable outputting detailed genotype likelihood information. Chromosome X TRs were genotyped using GangSTR v2.4.4 with additional options --bam-samps and --samp-sex to interpret sample sex for chromosome X. A separate GangSTR job was run for each family on each chromosome resulting in separate VCF files for each.

Genotypes were then subject to call-level filtering using dumpSTR, which is included in the TRTools toolkit v1.0.0^35^. DumpSTR was applied separately to each VCF with parameters --min-call-DP 20 --max-call-DP 1000 --filter-spanbound-only --filter-badCI --require-support 2 --readlen 150. Male chromosome X genotypes were filtered separately using the same parameters except with --min-call-DP 10. These options remove genotypes with too low or too high coverage, with only spanning or flanking reads identified indicating poor alignment, and with maximum likelihood genotypes falling outside 95% confidence intervals reported by GangSTR. After call-level filtering, each sample was examined for call-level missingness. All samples had >90% call rate and no outliers were identified.

Filtered VCFs from each phase were then merged using mergeSTR (TRTools v1.0.0) with default parameters. The merged VCF was then used as input to dumpSTR to compute locus-level filters using the parameters --min-locus-hwep 10^-5^ --min-locus-callrate 0.8 --filter-regions GRCh38GenomicSuperDup.sorted.gz --filter-regions-names SEGDUP to remove genotypes overlapping segmental duplications. The file GRCh38GenomicSuperDup.sorted.gz was obtained using the UCSC Table Browser ^36^ (hg38.genomicSuperDups table). For chromosome X, the Hardy-Weinberg Equilibrium filter was applied only to females. Filters obtained from analyzing each phase were combined and any TRs failing locus-level filters in any phase were removed from further analysis.

### Identifying de novo TR mutations

We developed a method, MonSTR (https://github.com/gymreklab/STRDenovoTools/), to identify *de novo* TR mutations from genome-wide TR genotypes obtained from GangSTR or HipSTR^37^. Our method extends code originally included in the HipSTR software (https://github.com/tfwillems/HipSTR). MonSTR is a model-based method that evaluates the joint likelihood of all genotypes of each parent-offspring trio and outputs a posterior estimate of a mutation occurring at each TR in each child. A full description of the MonSTR method is given in the **Supplementary Methods**.

MonSTR v1.0.0 was called separately on each family after applying call-level and locus-level genotype filters described above. MonSTR was called with non-default parameters --max-num-alleles 100 --include-invariant --gangstr --require-all-children --output-all-loci --min-num-encl-child 3 --max-perc-encl-parent 0.05 --min-encl-match 0.9 --min-total-encl 10 --posterior-threshold 0.5. Autosomes were run with the --default-prior −3 and chromosome X was run with the --naive option. These options remove TRs with too many alleles which are more likely to be error-prone, process all TRs even if no variation was observed, indicate to use GangSTR-output likelihoods (rather than HipSTR), only output loci if both children in the quad were analyzed, output all loci even if no mutation was observed, apply a constant prior of per-locus mutation rate of 10^-3^, require *de novo* mutation alleles to be supported by at least 3 enclosing reads, require *de novo* mutation alleles to be supported by fewer than 5% of parent enclosing reads, require 90% of enclosing reads in each sample to match the genotype call, require a minimum of 10 enclosing reads per sample in the family, and label calls with posterior probability ≥0.5 as mutations.

Resulting mutation lists output by MonSTR were subject to further quality control. We filtered families with likely sample contamination evidenced by extreme mutation counts (7 families, number of mutations > 1000), outlier mutation rates (16 families with number of mutations < 20 and > 241), mutations for which both children in the family were identified as having mutations at the same TR (n=43,239), and TRs with more than 25 mutations identified (n=15) as these are likely error-prone loci. We further filtered: calls for which the child was homozygous for the new allele (n=214,639), loci with a strong bias toward only observing contractions or expansions (n=179, two-sided binomial p<0.0001). We initially observed that mutations for which the parent of origin was homozygous often appeared to be erroneous due to drop out of one allele at heterozygous parents. This was most apparent for large mutations (± ≥ 5 repeat units) involving longer alleles difficult to span with short reads. We thus further required the new alleles to be supported by at least 6 enclosing reads in the child when the parent was called as homozygous.

Our stringent filtering of input genotypes and resulting mutations is unlikely to capture large repeat expansions, which are often not supported by enclosing reads because the resulting alleles are longer than Illumina read lengths. Thus, genotype likelihoods are more spread out and posterior estimates at these loci are lower and they will fail many of the QC options specified above. To additionally identify candidate expansions, we called MonSTR again on each family using the non-default parameter --naive-expansions-frr 3,8 which looks for TRs for which either: (1) the child has at least three fully repetitive reads and both parents have none or (2) the child has at least 8 flanking reads supporting an allele longer than any allele supported in either parent. We filtered candidate expansions identified in more than 3 samples, as we expect expansions to be rare. A total of 78 candidate expansions were identified across all families (**Supplementary Table 8**). These were merged with the total list of mutations for downstream analysis.

### Evaluating MonSTR on simulated WGS data

We created 78 quad families with 100 TR loci randomly selected from 1,024,318 genome-wide TRs passing all filters described above in the SSC cohort. One simulated quad family consists of the father, mother, child with known mutation (proband), and child with no mutation (control). We tested the ability of our entire pipeline to genotype TRs with GangSTR and call *de novo* mutations with MonSTR. To test the effect of depth of coverage, we generated data sets with 1-50x mean coverage with a mutation size of +1 or −1 repeat unit changes in the proband. To test the effect of TR mutation size, we generated WGS data with 40x coverage and known mutations in probands ranging from −10 to 30 repeat units step size changes. Contraction mutations that would have resulted in negative repeat copy numbers were excluded. For both tests, we simulated data under three scenarios: (1) both parents with homozygous reference TR genotypes, (2) one parent heterozygous, (3) both parents heterozygous (Extended Data Fig. 1).

WGS data were simulated using ART_illumina^38^ v2.5.8 with non-default parameters -ss HS25 (HiSeq 2500 simulation profile), -l 150 (150b reads), -p (paired-end reads), -f coverage (coverage was set as described above), -m 500 (mean fragment size) and -s 100 (standard deviation of fragment size). ART_illumina was applied to fasta files generated from 10Kb windows surrounding each TR locus, applying any mutations as described above. The resulting fastq files were aligned to the hg38 reference genome using bwa mem^39^ v0.7.12-r1039 with non-default parameter -R “@RG\tID:sample_id\tSM:sample_id”, which sets the read group tag ID and sample name to sample_id for each simulated sample. TRs were genotyped from aligned reads jointly across all members of the same family with GangSTR using identical settings to those applied to SSC data.

We tested three mutation calling settings: a naïve mutation calling method based on hard genotype calls, MonSTR using default parameters, and MonSTR using an identical set of filters as applied to SSC data. We found overall all methods perform similarly well above 30x coverage. At lower coverage, MonSTR’s model-based method achieves reduced sensitivity but greater specificity compared to a naïve mutation calling pipeline (Extended Data Fig. 1).

### Comparison to previously reported mutation rates

Mutation rates for CODIS markers were obtained from the National Institute of Standards and Technology (NIST) website (https://strbase.nist.gov/mutation.htm). 95% confidence intervals on the estimated number of mutations that should be observed in SSC were obtained by drawing mutation counts from a binomial distribution with n=the total number of children genotyped at each locus and p=the NIST estimated mutation rate (Extended Data Fig. 2c). Intervals were obtained based on 1,000 simulations.

Genome-wide autosomal TR mutation rates and constraint scores estimated using MUTEA^12^ were obtained from https://s3-us-west-2.amazonaws.com/strconstraint/Gymrek_etal_SupplementalData1_v2.bed.gz (columns est_logmu_ml and zscore_2). TRs were converted from hg19 to hg38 coordinates using the liftOver tool available from the UCSC Genome Browser Store free for academic use (https://genome-store.ucsc.edu/). We intersected the lifted over coordinates with the GangSTR reference using the intersectBed tool included in BEDTools v2.28.0^40^. Only TRs overlapping GangSTR TRs by at least 50% (−f 0.5) and with the same repeat unit length in each set were used for analysis.

### Evaluation of mutations with capillary electrophoresis (CE) fragment analysis

Whole blood-derived genomic DNA for 5 SSC quad families was obtained through SFARI Base to validate a subset of TR mutation calls. For each candidate TR, we designed primers to amplify the TR and surrounding region (**Supplementary Table 3**). A universal M13(−21) sequence (5’-TGTAAAACGACGGCCAGT-3’) was appended to each forward primer. We then amplified each TR using a three-primer reaction previously described^41^ consisting of the forward primer with the M13(−21) sequence, the reverse primer, and a third primer consisting of the M13(−21) sequence labeled with a fluorophore.

The forward (with M13(−21) sequence) and reverse primers for each TR were purchased through IDT. The labeled M13 primers were obtained through ThermoFisher (#450007) with fluorescent labels added to the 5’ ends (either FAM, VIC, NED, or PET). TRs were amplified using the forward and reverse primers plus an M13 primer with one of the four fluorophores with GoTaq polymerase (Promega #PRM7123) using PCR program: 94°C for 5 minutes, followed by 30 cycles of 94°C for 30 seconds, 58°C for 45 seconds, 72°C for 45 seconds, followed by 8 cycles of 94°C for 30 seconds, 53°C for 45 seconds, 72°C for 45 seconds, followed by 72°C for 30 minutes.

The CGG repeat at chr7:103989357 in the 5’UTR of *RELN* could not be amplified using the three-primer method and was genotyped using published primers^42^ (forward: 5′-FAMCGCCTTCTTCTCGCCTTCTC-3′ and reverse: 5′-CGAAAAGCGGGGGTAATAGC-3′). The TR was amplified with HotStarTaq Polymerase (Qiagen #203203) using PCR program: 95°C for 15 minutes, followed by 35 cycles of 94°C for 45 seconds, 58°C for 60 seconds, 72°C for 60 seconds, followed by 72°C for 30 minutes.

Fragment analysis of PCR products was performed on a ThermoFisher SeqStudio instrument using the GSLIZ1200 ladder, G5 (DS-33) dye set, and long fragment analysis options. Resulting .fsa files were analyzed by manual review in GeneMapper (ThermoFisher # 4475073).

### Analysis of mutation directionality bias

The observed bias of longer alleles to contract and shorter alleles to expand (Fig. 2c) could potentially be explained by genotyping errors at heterozygous loci due to “heterozygote dropout” of long alleles, leading to erroneous homozygous genotype calls. To reduce the potential impact of heterozygote dropout on apparent mutation directionality, we restricted this analysis to mutations with an absolute size of ≤5 units. When analyzing mutations from heterozygous vs. homozygous parents (Extended Data Fig. 3d-e), we further restricted to mutations consisting of a single unit and for which the child had at least 10 enclosing reads supporting the *de novo* allele, indicating the allele could be easily spanned and would be less prone to dropout.

### Determinants of TR mutation rates

Genomic and epigenomic features for each TR were compiled from a variety of resources. BEDTools intersect v2.28.0 was used to overlap GangSTR reference TRs for each annotation after using liftOver to convert each to hg38 coordinates. Recombination rates^43^ were obtained from https://github.com/cbherer/Bherer_etal_SexualDimorphismRecombination/blob/master/Refined_genetic_map_b37.tar.gz. Rates were log_10_ transformed with a pseudo count of 0.0001 to avoid infinite values. TRs with scaled recombination rates >-3 were filtered. PhastCons^44^ annotations were obtained from http://hgdownload.cse.ucsc.edu/goldenpath/hg38/phastCons100way/. Values were log_10_ transformed with a pseudo count of 0.0001 to avoid infinite values. TRs with transformed scores >-4 were filtered. Nucleosome occupancy scores were obtained from http://hgdownload.soe.ucsc.edu/goldenPath/hg18/database/uwNucOccMec.txt.gz (Mec^45^) and http://hgdownload.soe.ucsc.edu/goldenPath/hg18/database/uwNucOccDennis.txt.gz (Dennis^46,47^). Notably these two annotations were scored in opposite directions and should be anti-correlated. TRs with annotations >-4 or <2, respectively, were filtered. Conserved promoter annotations were obtained from Table S9 of An *et al*^11^. DNAseI hypersensitivity peaks based on 125 cell types profiled by the ENCODE Project^48^ were obtained from the UCSC genome browser (http://hgdownload.cse.ucsc.edu/goldenpath/hg19/encodeDCC/wgEncodeRegDnaseClustered/) and were treated as binary features. Histone modification peaks for embryonic stem cells (H1) were obtained from the Encode Project website (https://www.encodeproject.org) and were treated as binary features (accessions ENCFF180RPI, ENCFF720LVE, ENCFF835TGA, ENCFF219TGT, ENCFF483GVK, ENCFF695ZZV, ENCFF781GRI, ENCFF067WBB, ENCFF714VTU, ENCFF073WSF). GC content for TR motifs and for varying size windows around each TR were computed using a custom script based on the hg38 reference genome.

We fit a Poisson regression model to predict mutation counts in unaffected individuals at each TR based on these features. A separate model was fit for each feature and each repeat unit length, in each case using TR reference length (in bp) as a covariate. In each model the exposure was set to the number of observed transmissions in unaffected individuals. Models were fit using the discrete.discrete_model.Poisson module from the Python statsmodels library v0.10.1 (https://www.statsmodels.org/).

### Mutation burden statistical testing

Mutation excess in probands vs. non-ASD siblings was tested using a paired t-test as implemented in the Python scipy library v1.3.1 (https://docs.scipy.org/doc/scipy/reference/index.html) function scipy.stats.ttest_rel. We compared a vector of counts of mutations in probands to a vector of counts in mutations in non-ASD siblings, ordered by family ID.

Comparison of TR mutation burden in probands vs non-ASD siblings was also computed after adjusting for the father’s age at birth. We used the Python statsmodels ordinary least squares regression module to regress unaffected mutation counts on paternal age. We then used this model to compute residual mutation counts in each sample after regressing on paternal age.

Relative risk was computed as the ratio of the mean number of mutations in probands vs. non-ASD siblings. Relative risk of greater than 1 indicates a higher burden in the probands. We estimated a 95% confidence interval on the fraction of mutations 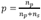 in each category that are in probands vs. siblings based on a binomial distribution 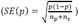 where *n_p_* and *n_s_* are the number of mutations observed in probands and siblings, respectively. We then used the upper and lower bounds on the fraction of mutations in probands *p_low_* = *p* – 1.96*SE(p)*; *p_high_* = *p* + 1.96*SE(p)* to compute the corresponding 95% confidence intervals for relative risk as 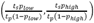 where *t_s_* and *t_p_* are the total number of sibling and proband samples considered, respectively. Gene annotations were obtained from the UCSC Table Browser^36^ using the hg38 reference genome. Fetal brain promoter and enhancer annotations were obtained from fetal brain male ChromHMM^49^ annotations available on the ENCODE Project website (https://www.encodeproject.org/; accession ENCSR770CMJ).

The contribution of *de novo* TR mutations to ASD risk was calculated by taking the difference in total autosomal mutations identified in probands vs. siblings divided by the number of probands, as was done in a previous study of non-coding mutations in ASD^50^.

### Enrichment of common variant risk

GWAS SNP associations were downloaded from GWAS catalog^51^ (ASD [EFO_0003756] n= 637 SNPs; SCZ [EFO_0000692] n= 3,476; EA [EFO_0004784] n=3,966). We tested whether TR mutations falling within 50kb of GWAS SNPs for each trait showed increased burden in probands vs. siblings by performing a Mann Whitney test (Python function scipy.stats.mannwhitneyu) comparing mutation counts in probands vs. non-ASD siblings. We performed an additional test excluding mutations resulting in alleles with AF<0.05.

### Gene expression analysis

The Developmental Transcriptome dataset containing RNA-seq normalized gene expression values and meta-data for developmental brain tissue regions was downloaded from the BrainSpan Atlas of the Developing Human Brain^52^ (https://www.brainspan.org/static/download.html). Expression values were log-transformed before analysis, adding a pseudo count of 0.01 to avoid 0 values. We excluded brain structures “CB”, “LGE”, “CGE”, “URL”, “DTH”, “M1C-S1c”, “Ocx”, “MGE”, “PCx”, and “TCx” since those structures only had data for male samples at 3 or fewer time points. We used a one-sided Mann Whitney test (scipy.stats.mannwhitneyu) to compare the distribution of expression in genes with only proband mutations vs. genes with only unaffected sibling mutations separately for each tissue. Meta-analysis across all 32 brain regions was performed using Fisher’s method to combine p-values. Expression STR summary statistics were obtained from Supplementary Data 2 of Fotsing, *et al*.^53^.

### Inferring selection coefficients using SISTR

We developed SISTR (Selection Inference at short TRs), a population genetics framework for inferring selection coefficients at individual TR loci. SISTR fits an evolutionary model of TR variation that includes mutation, genetic drift, and negative natural selection to available empirical allele frequencies to infer the posterior distribution of selection coefficients. Our mutation model is based on a modified version of the generalized stepwise mutation model (GSM)^54^. To model negative selection, we assume the central allele at each TR has optimal fitness (*w*=1), and that the fitness of other alleles is based on their difference in size from the optimal allele.

SISTR applies approximate Bayesian computation (ABC) based on a previously described forward simulation technique^55^ to infer per-locus selection coefficients by fitting allele frequencies for one TR at a time given a predefined optimal allele length and fixed set of mutation parameters. Our method outputs the median posterior estimate of *s* and computes a likelihood ratio test comparing the likelihood of the inferred *s* value to the likelihood of *s*=0. Full descriptions of the mutation and selection models and the SISTR inference method are given in the **Supplementary Methods**.

For each TR with a repeat unit length of 2-4bp, we used SISTR to estimate selection coefficients based on allele frequencies in SSC parents. We set the optimal allele length at each TR to the modal allele and used mutation parameters described in the **Supplementary Methods** as input. We excluded TRs with repeat lengths in hg38 <11 units for dinucleotides, <5 units for trinucleotides, and <7 repeats for tetranucleotides, since those repeats are typically not polymorphic. We included only TRs for which precise copy numbers could be inferred in at least 80% of SSC parents. After filtering, 82,223 STRs remained for analysis.

We used the Benjamini-Hochberg procedure^56^ to adjust p-values for multiple hypothesis testing. To identify TRs under significant selection, we chose TRs with adjusted p-value <0.01, corresponding to a false discovery rate of 1%. Allele-specific selection coefficients, which can be interpreted as pathogenicity scores, were computed as (*a − opt*)*s*, where *a* is the number of repeat copies for the *de novo* allele, *opt* is the optimum (modal) repeat and *s* is the selection coefficient for the TR inferred using SISTR.

Gene-level constraint metrics (pLI and missense Z score) were obtained from https://storage.googleapis.com/gnomadpublic/release/2.1.1/constraint/gnomad.v2.1.1.lof_metrics.by_gene.txt.bgz.

## Acknowledgments

This study was supported by the Simons Foundation Autism Research Initiative (SFARI Grant #630705). I.M. was supported by a predoctoral fellowship from the Autism Science Foundation. B.H. and M.G. were supported in part by the Office Of The Director, National Institutes of Health under Award Number DP5OD024577. M.G. was additionally supported in part by NIH/NHGRI grants R01HG010149 and R21HG010070, and NIH/NIMH grant R01 MH113715. K.E.L. was supported by the National Institutes of Health grant R35GM119856. The authors thank Joseph Gleeson, Jonathan Sebat, Abraham Palmer, and Alon Goren for helpful comments on this study.

## Author contributions

I.M. performed TR genotyping, identification of *de novo* mutations, and downstream analyses and helped write the manuscript. B.H. developed SISTR and performed analysis of TR selection scores in the SSC cohort. Nima M. helped design GangSTR analysis and filtering settings and analyses to evaluate MonSTR. Nichole M. performed capillary electrophoresis validation experiments. M.L. performed TR annotation for identification of determinants of TR mutation rates. R.Y. designed AWS cloud analysis pipelines. S.S.-B. helped design and set up validation experiments. K.E.L. conceived the SISTR method, supervised analysis of TR selection scores, and drafted the manuscript. M.G. conceived the study, designed and performed analyses, and drafted the manuscript. All authors have read and approved the final manuscript.

## Data Availability

Upon acceptance of this study for publication, all TR genotypes and mutation calls will be deposited at SFARI Base. Per-locus selection scores computed by SISTR are provided in **Supplementary Data 1**.

## Code Availability

The (1) MonSTR software for identifying TR mutations and (2) SISTR software for prioritizing TR mutations are open source and available on Github: (1) https://github.com/gymreklab/STRDenovoTools and (2) https://github.com/BonnieCSE/SISTR. The code used to generate figures and results for this study are available at https://github.com/gymreklab/ssc-denovos-paper.

## Competing interests

The authors have no competing financial interests to disclose.

---

**Extended Data Figure 1:**
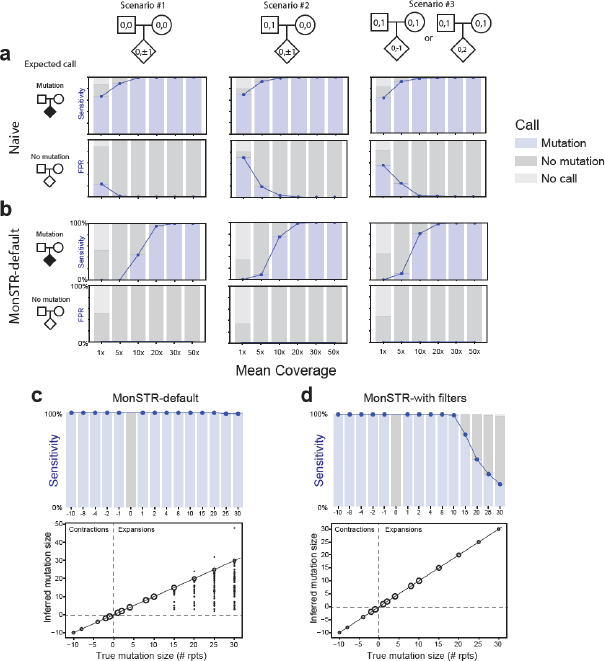
Evaluation of MonSTR using simulated data. **a. Evaluation of a naïve TR mutation calling method**. WGS was simulated for probands with mutations and controls with no mutation under three different scenarios for a range of mean sequencing coverages. Top plots show the sensitivity (blue line). Bottom plots show the false positive rate (FPR). Shaded bars show the percent of transmissions called as mutation (blue), no mutation (dark gray), or no call (light ray). **b. Evaluation of MonSTR’s default model-based method**. Plots are the same as in **a**. but based on MonSTR’s default model. Note FPR lines are not visible because all are at 0%. **c. Evaluation of TR mutation calling using default MonSTR settings as a function of mutation size**. The top plot is the same as in **a-b**, and shows the sensitivity to detect mutations as a function of their size. The bottom plot compares the estimated called mutation size (y-axis) compared to the true simulated mutation size (x-axis). Bubble sizes represent the number of mutation calls represented at each point. **d. Evaluation of TR mutation as a function of mutation size after quality filtering**. Plots are same as in **c**, but using the stringent quality filters in MonSTR applied to analyze the SSC cohort. Compared to default settings, sensitivity is decreased especially for larger expansions but inferred mutation sizes are unbiased. All plots are based on simulation of 100 randomly chosen TR loci (**Methods**). **c-d** show results for scenario #1.

**Extended Data Figure 2:**
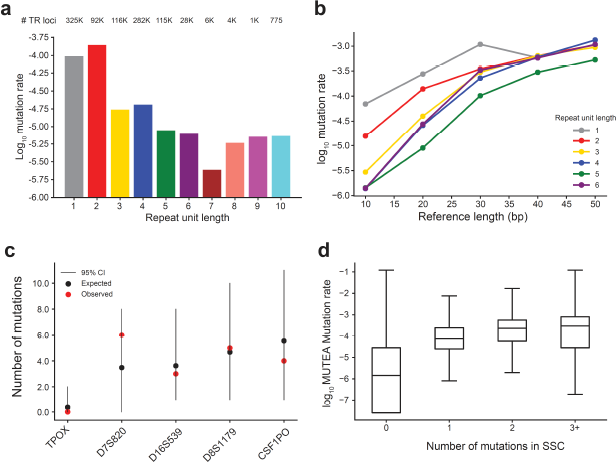
Patterns of TR mutation recapitulate previously observed trends. **a. Distribution of average TR mutation rates by period**. For each repeat unit length (x-axis), bars give the genome-wide estimated TR mutation rate (y-axis, log_10_ scale). Average mutation rates were computed as the total number of mutations divided by the total number of observed parent to child transmissions observed and passing all quality filters. The total numbers of TRs considered in each category are annotated (top). **b. TR Mutation rate vs. length**. The x-axis shows the TR reference length (in base pairs) and the y-axis shows the mutation rate (log_10_ scale) estimated across all TRs with each reference length within each category (repeat unit length). Colors denote different repeat unit lengths (gray=homopolymers; red=dinucleotides; gold=trinucleotides; blue=tetranucleotides; green=pentanucleotides; purple=hexanucleotides). **c. Number of *de novo* TR mutations observed (red) for CODIS markers**. Black dots show expected mutation counts and lines give 95% confidence intervals based on mutation rates reported by NIST (https://strbase.nist.gov/mutation.htm). Each x-axis category denotes a separate CODIS marker. **d. Observed TR mutation counts concordant with MUTEA**. Boxes show the distribution of log_10_ mutation rates estimated by MUTEA^12^ (y-axis) at each TR with a given number of mutations observed in SSC (x-axis). Black middle lines give medians and boxes span from the 25th percentile (Q1) to the 75th percentile (Q3). Whiskers extend to Q1-1.5*IQR (bottom) and Q3+1.5*IQR (top), where IQR gives the interquartile range (Q3-Q1).

**Extended Data Figure 3:**
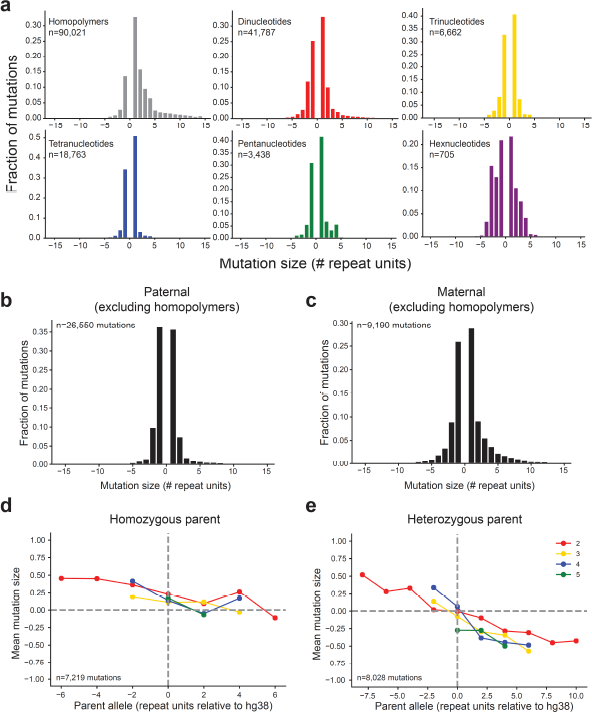
Biases in TR mutation sizes. **a. Mutation step size distributions by period (repeat unit length)**. Histograms show the distribution (y-axis, fraction of total) of *de novo* TR mutation sizes (x-axis, number of repeat units) for each repeat unit length. Mutations <0 denote contractions and >0 denote expansions. Colors denote different repeat unit lengths (gray=homopolymers; red=dinucleotides; gold=trinucleotides; blue=tetranucleotides; green=pentanucleotides; purple=hexanucleotides). **b-c. Mutation step size distributions by parental origin**. Histograms show the distribution of *de novo* TR mutation sizes for mutations arising in the paternal (**b**) and maternal (**c**) germlines (homopolymers excluded). **d-e. Mutation directionality bias in homozygous vs. heterozygous parents**. In each plot, the x-axis gives the size of the parent allele relative to the reference genome (hg38). The y-axis gives the mean mutation size in terms of number of repeat units across all mutations with a given parent allele length. A separate colored line is shown for each repeat unit length (red=dinucleotides; gold=trinucleotides; blue=tetranucleotides; green=pentanucleotides). Plots are restricted to mutations that were successfully phased to either the mother or the father for which the parent of origin was homozygous (**b**) or heterozygous (**c**). To restrict to highest confidence mutations, these plots are based only on mutations with step size of ±1 and for which the child had more than 10 enclosing reads supporting the *de novo* allele.

**Extended Data Figure 4:**
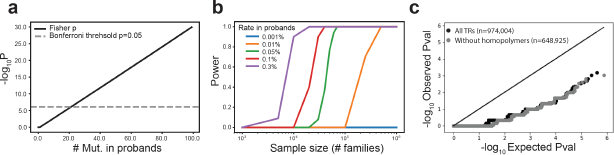
Power to detect per-locus TR mutation enrichments. **a. Number of recurrent mutations required to reach genome-wide significance**. We performed a Fisher’s exact test to test for an excess of mutations in probands vs. non-ASD siblings, for a different number of hypothetical mutation counts in probands (x-axis) and assuming 0 mutations observed in non-ASD siblings. The black line shows the two-sided p-value obtained for each test. The gray dashed line denotes the p-value required to meet a genome-wide significance of p<0.05 with Bonferroni multiple testing correction. Genome-wide significance can only be obtained in the SSC cohort if a TR has 20 or more mutations in probands and zero in non-ASD siblings. **b. Sample sizes required to identify genome-wide significant TRs**. The x-axis shows sample size (log_10_ scale) in terms of the number of quad families analyzed. Each line represents a different rate of mutation at a particular TR in probands, assuming 0 mutations at that TR in siblings (blue=0.001%; orange=0.01%; green=0.05%; red=0.1%; purple=0.3%). The y-axis shows the power to detect a specific TR at genome-wide significance for each rate. **c. Quantile-Quantile plots for per-locus TR mutation burden testing**. For each TR we performed a Fisher’s exact test to test for an excess of mutations in probands vs. siblings. The x-axis gives expected −log_10_ p-values under a null (uniform) distribution. The y-axis gives observed −log_10_ p-values from burden tests. Each dot represents a single TR. Black=all TRs. Gray=homopolymers excluded.

**Extended Data Figure 5:**
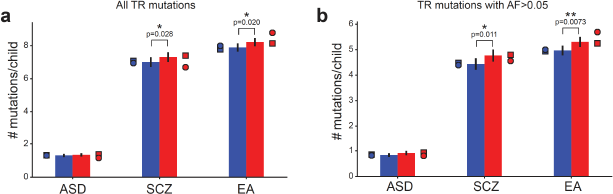
TR mutation burden near GWAS loci for ASD and related traits. Bars show mean mutation counts in probands (red) vs. non-ASD siblings (blue) considering all TRs (**a**) or TRs resulting in alleles with frequency >5% in controls (**b**) that are within 10kb of published GWAS SNPs (ASD=autism spectrum disorder; SCZ=schizophrenia; EA=educational attainment). Error bars give 95% confidence intervals. Single asterisks denote nominally significant increases (p<0.05). Double asterisks denote trends that are significant after Bonferroni correction for the six categories tested. Circles and squares show counts for females and males, respectively.

**Extended Data Figure 6:**
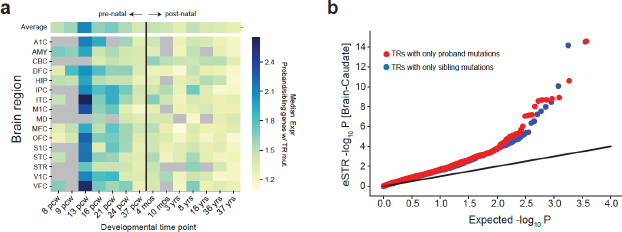
Proband *de novo* TR mutations enriched in brain-expressed genes. **a. Ratio of median expression in proband-only genes to control-only genes across time points**. The heatmap shows the ratio of the median expression of genes with only proband mutations to that of genes with only mutations in non-ASD. Each row shows a different brain structure from the BrainSpan dataset. Each column shows a different developmental timepoint. The black vertical line separates prenatal from postnatal time points. Gray boxes indicate no data was available for that time point. Brain structure acronyms are the same as in Fig. 2. **b. Proband TR mutations enriched for brain expression STRs**. The quantile-quantile plot shows the distribution of expression STR (eSTR) p-values based on associating TR length with gene expression in Brain-Caudate samples in the GTEx cohort^59^. Each point represents a TR by gene association test using a linear regression model^53^. The x-axis gives expected −log_10_ p-values and the y-axis gives observed −log_10_ p-values. Red points show TRs with at least one *de novo* mutation in probands and 0 in controls. Blue points show TRs with at least one *de novo* mutation in controls and 0 in probands. We found no significant difference in either Brain-Cerebellum or the other 15 non-brain tissues analyzed in that study, which expected should not be relevant to ASD (not shown).

**Extended Data Figure 7:**
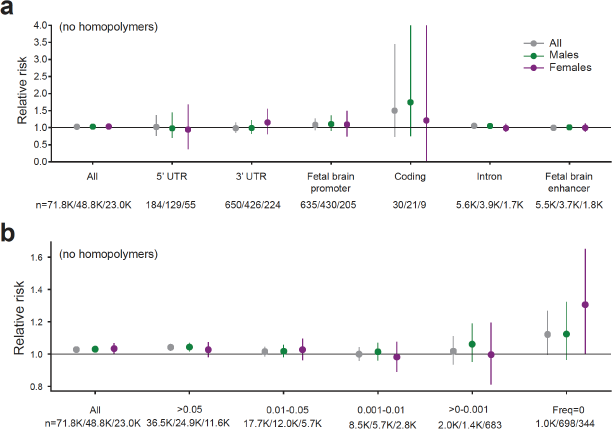
TR mutation burden in ASD excluding homopolymers. **a. Mutation burden by gene annotation**. Dots show estimated relative risk and lines give 95% confidence intervals. Gray=all samples; green=males only; purple=females only. **b. Mutation burden by frequency of the allele arising by *de novo* mutation**. The x-axis stratifies mutations based on non-overlapping bins of the frequency of the *de novo* allele in healthy controls (SSC parents). “All” includes all mutations. For other allele frequency bins, only TRs for which precise copy numbers could be inferred in at least 80% of SSC parents are included. The y-axis gives RR in probands vs. non-ASD siblings. AF=allele frequency. Both plots show only TRs with repeat unit length >1bp.

**Extended Data Figure 8:**
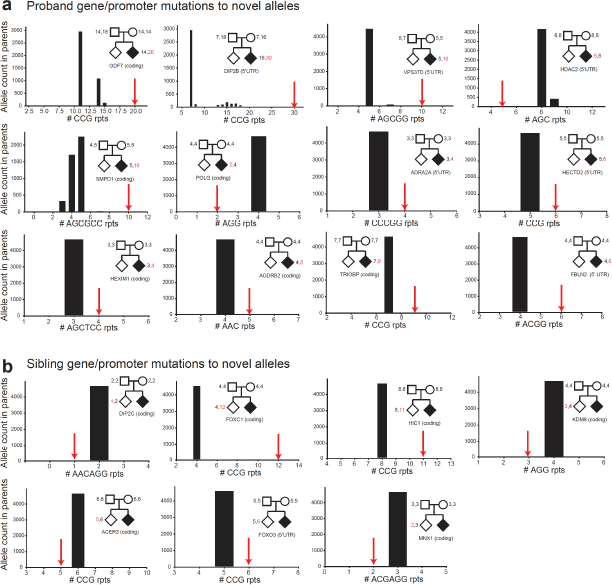
Gene and 5’UTR mutations to novel alleles. **a. Mutations in probands at coding or 5’UTR TRs to unobserved alleles**. Each panel shows a *de novo* TR mutation observed in ASD probands to an allele (x-axis, repeat copy number) not observed in SSC parents. Black histograms give the allele counts in parents. Red arrows denote the allele resulting from each example *de novo* TR mutation. Pedigrees show genotypes of parents and the child with the mutation, where ASD probands are diamond shaped colored in black and non-ASD siblings are uncolored diamond. The text below pedigrees gives the gene and region in which the mutation occurred. **b. Mutations in non-ASD siblings at coding or 5’UTR TRs to unobserved alleles**. Plots are the same as in **a**.

**Extended Data Figure 9:**
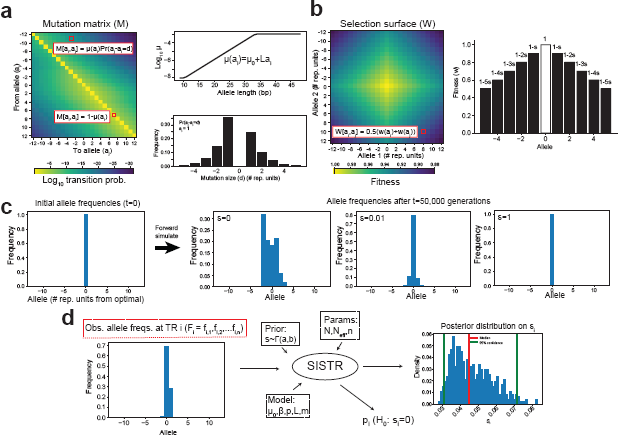
A method to estimate selection coefficients for STRs. **a. Short TR mutation model**. Mutation is modeled by a stochastic mutation matrix with length-dependent mutation rates and mutation sizes following a geometric distribution with a directional bias toward the central allele. Unless otherwise indicated, alleles are specified in terms of the number of repeat units away from the central, or modal, allele at each STR. **b. STR selection model**. Negative selection is modeled by a diploid selection surface constructed as a function of the fitness of the individual alleles. The fitness of each allele is calculated as a function of a selection coefficient *s*, where the central allele has optimal fitness (*w*=1), and the fitness of other alleles is a function of the number of repeat units away from the optimal allele. **c. Example output of forward simulations of allele frequencies**. The simulation starts with one ancestral (“optimal”) allele. As *s* increases, variability in the resulting allele frequency distributions decreases as the less fit alleles are removed by natural selection. **d. Overview of per-STR selection inference using Approximate Bayesian Computation**. For each STR, the method takes a prior on *s*, mutation model, and demographic parameters, and the observed allele frequency distribution as input. It outputs a posterior distribution of *s* and a p-value from a likelihood ratio test of whether a model with selection fits better than a model without selection (*s*=0).

**Extended Data Figure 10:**
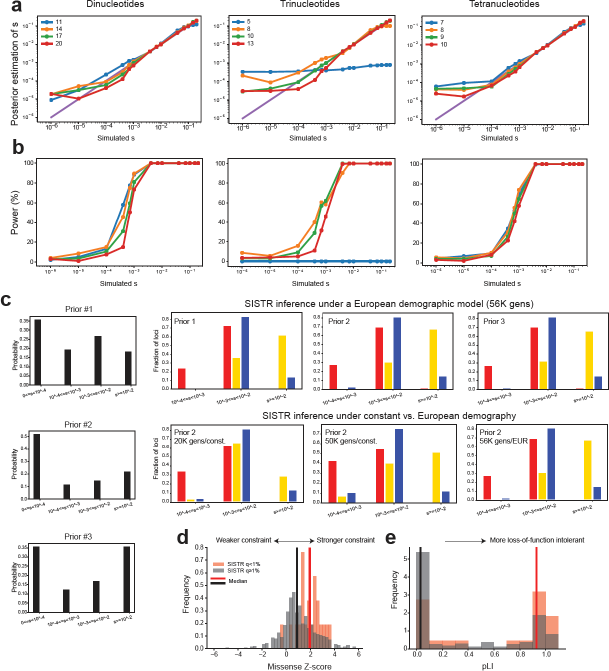
Evaluation of SISTR. **a. Comparison of true vs. inferred per-locus selection coefficients**. The x-axis shows the true simulated value of *s*, and the y-axis shows the mean *s* value inferred by SISTR across 200 simulation replicates. **b. Power to detect negative selection as a function of *s***. The x-axis shows the true simulated value of *s*, and the y-axis gives the power to reject the null hypothesis that *s*=0. Left, middle, and right panels show results using models for dinucleotide, trinucleotide, and tetranucleotide TRs, respectively. **c. Inferred genome-wide distribution of *s* is robust to prior choice and demographic models**. We applied SISTR genome-wide using 2 different demographic models (**Supplementary Methods**) and 3 different prior distributions (left panels) on *s*. Right panels show the inferred genome-wide distribution of *s* using different combinations of priors and demographic models. Red, yellow, and blue denote dinucleotides, trinucleotides, and tetranucleotides, respectively. **d. Genes containing coding STRs under strong selection are more missense-constrained**. The x-axis gives the missense constraint Z-score reported by Gnomad^33^. The y-axis gives the frequency of genes with each missense Z-score. **e. Genes containing coding STRs under strong selection are more loss-of-function intolerant**. The x-axis gives the pLI score measuring loss of function intolerance of each gene reported by Gnomad. For **d** and **e**, black bars show the distribution for all genes containing an STR not inferred to be under selection (adjusted p ≥1%) and red bars show the distribution for all genes containing an STR inferred to be under selection (adjusted p<1%). Vertical lines show medians of each distribution.

## Notes

https://github.com/gymreklab/ssc-denovos-paper

### Competing Interest Statement

The authors have declared no competing interest.

