## Supplemental Information for "Genome-wide patterns of *de novo* tandem repeat mutations and their contribution to autism spectrum disorders"

### Supplementary Information

### Supplementary Methods

#### *MonSTR model and method details*

MonSTR is a method to identify *de novo* TR mutations from genotypes obtained from whole genome sequencing (WGS). It takes genotype likelihoods as input and outputs the estimated posterior probability of a mutation occurring at each TR in each child. The MonSTR implementation extends code originally included in the HipSTR[1] software (https://github.com/tfwillems/HipSTR) written by Thomas Willems. The full MonSTR package is available at https://github.com/gymreklab/STRDenovoTools/.

**MonSTR statistical model**

MonSTR's likelihood model is based on a previously published model for identifying *de novo* point mutations[1]. It considers a single trio (father, mother, and child) at a time. Let $G_m = (g_{m1}, g_{m2})$ be the diploid genotype (allele lengths) of the mother, $G_f = (g_{f1}, g_{f2})$ the diploid genotype of the father, and $G_c = (g_{c1}, g_{c2})$ the diploid genotype of the child in a trio. Here we will assume that the child's genotype is ordered, with allele $g_{c1}$ derived from the mother and $g_{c2}$ from the father. The likelihood of a particular genotype configuration with no mutation can be defined as:

$$L(G_m, G_f, G_c \mid D) \propto M(G_m|D)F(G_f|D)C(G_c|D) \times T(G_c|\, G_m, G_f) \times P(G_m, G_f)$$

Where $D$ denotes available WGS data for the family at that locus, $M(G_m|D)$ denotes the likelihood of maternal genotype $G_m$ given the data, $F(G_f|D)$ denotes the likelihood of paternal genotype $G_f$ given the data, $C(G_f|D)$ denotes the likelihood of child genotype $f$ given the data, $T(G_c|\, G_m, G_f)$ is a transition probability, and $P(G_m, G_f)$ is the prior probability of observing the parent genotypes. Genotype likelihoods $M$, $F$, and $C$ are obtained directly from the GGL (genotype likelihood) field output by GangSTR. MonSTR is also compatible with the GL field (genotype likelihood) output by HipSTR. We assume a uniform prior on all genotypes so the prior term is dropped. For simplicity below we drop the $D$ term.

The likelihood the observed data with no mutation is computed as:

$$L(nomut) = \sum_{G_m} \sum_{G_f} M(G_m)F(G_f) \sum_{G_c \in I} T(G_c|\, G_m, G_f) \times C(G_c) \qquad \text{(Eq. 1)}$$

Where $I$ represents genotypes that could result from all possible transmission scenarios, i.e. $G_c = (g_{m1}, g_{f1})$, $(g_{m1}, g_{f2})$, $(g_{m2}, g_{f1})$, or $(g_{m2}, g_{f2})$. The transmission probability $T(G_c|\, G_m, G_f)$ is simply equal to 0.25 for any particular transmission.

The likelihood of a mutation occurring is computed as:

$$L(mut) = \sum_{G_m} \sum_{G_f} M(G_m)F(G_f) \sum_{G_c \in I} T(G_c|\, G_m, G_f) \left[\sum_k R(g_{c1}, k)C(g_{c1} + k, g_{c2}) + R(g_{c2}, k)C(g_{c1}, g_{c2} + k)\right] \qquad \text{(Eq. 2)}$$

Where $R(g, k)$ gives the transition probability of allele $g$ mutating by $k$ units. The above equation considers all four possible transition scenarios (similar to the likelihood for no mutation), followed by a mutation either to the maternally inherited allele ($g_{c1}$) or to the paternally inherited allele ($g_{c2}$). For this study we used a naive mutation transition probability treating each mutation size as equally likely. MonSTR also allows for specifying locus-specific mutation parameters (see below), which we did not find to significantly change our results.

Finally, MonSTR computes the posterior probability of a mutation as:

$$P(mut|D) = \frac{L(mut|D)P(mut)}{L(mut|D)P(mut) + L(nomut|D)(1 - P(mut))}$$

Where $P(mut)$ gives the prior probability of mutation.

**MonSTR statistical model for chromosome X**

The model presented above applies to autosomal TRs. MonSTR implements a modified version of the model for identifying mutations on chromosome X.

For chromosome X loci, males have haploid genotype calls. Thus, $G_f = (g_f)$ rather than $G_f = (g_{f1}, g_{f2})$. For female children, Equations 1 and 2 above have only slight modifications:

- $I$, which represents the set of child genotypes that could result from all possible transmission scenarios, includes $G_c = (g_{m1}, g_f)$ or $(g_{m2}, g_f)$.
- The transition probability $T(G_c|\, G_m, G_f)$ is set to 0.5 for each case.

For male children, $G_c = (g_c)$, and the following further modifications are made:

- $I$ includes child genotype possibilities $G_c = (g_{m1})$ or $(g_{m2})$.
- The transition probability only depends on the maternal genotype and $T(G_c|\, G_m)$ is set to 0.5 for each possible child genotype.
- Equation 2 (likelihood of mutation) is modified to only consider mutations from the mother:

$$L(mut) = \sum_{G_m} M(G_m) \sum_{G_c \in I} T(G_c|\, G_m) [\sum_k R(g_{c1}, k) C(g_{c1} + k)]$$

**Naïve mutation model**

In addition to the model-based mutation detection method described above, MonSTR also implements two naïve methods:

If the –naïve option is specified (see below for a full description of options), MonSTR will simply check if the child genotype call can be explained by Mendelian inheritance from genotypes called in the parents. Thus, it does not consider uncertainty from genotype likelihoods and outputs a binary result for mutation/no mutation rather than a posterior probability.

If the –naïve-expansions-frr <int1,int2> option is specified, MonSTR will output candidate expansion mutations that have either <int1> fully repetitive reads (FRRs; described previously[2,3]. Fully repetitive reads at a locus indicate a potential repeat expansion) in a child and 0 in parents, or <int2> flanking reads supporting an allele length greater than the largest allele observed in parents.

**Inferring parent of origin**

For each mutation identified, we attempted to infer the likely parent of origin based on the child and parent genotypes. Let $C = (c_1, c_2)$ be the child genotype, where $c_i$is the number of repeats called in the ith allele. Similarly, let $F = (f_1, f_2)$ and $M = (m_1, m_2)$ denote the father and mother genotypes, respectively. If $c_1 \in M$ and $c_1 \notin F$, we determine $c_2$ to be the new allele and phase the mutation to the father. If $c_2 \in M$ and $c_2 \notin F$, we determine $c_1$to be the new allele and phase the mutation to the father. Similarly, if $c_1 \in F$ and $c_1 \notin M$ or $c_2 \in F$ and $c_2 \notin M$, the new allele is determined to be $c_1$ or $c_2$, respectively, and the mutation is phased to the mother. All mutations not meeting the above conditions were labeled with phase as unknown.

**MonSTR implementation details**

MonSTR is implemented as a standalone command-line program written in C++. To improve speed and integration with other tools, it uses standard file formats and leverages the htslib library (www.htslib.org) and previously written VCF parsing libraries implemented in HipSTR[4].

MonSTR takes as input a multi-sample VCF with genotype likelihoods for each sample generated by HipSTR or GangSTR and a pedigree file (in plink[5] .fam format https://www.cog-genomics.org/plink2/formats#fam). It outputs a tab-delimited file describing mutation posterior probabilities, mutation sizes, and parent of origin inferences, for each trio at each TR.

MonSTR can call mutations using either “model-based” (default) or “naïve” mode (options –naïve or –naïve-expansions-frr described above). In the “model-based” option, users can choose either a global prior probability of mutation (default to $10^{-5}$ mutations per locus per generation) or input a file with per-locus mutation rates to use as priors. Further, the transition probabilities $R(g, k)$ can be set either to a uniform probability to mutate to any allele, or users may input values of parameters $\beta$ (length dependent direction bias), $p$

(mutation step size geometric distribution parameter), and the central allele, which are described in detail below, to implement more detailed step size distribution models.

MonSTR also offers many options to discard individual calls based on properties including their coverage, genotype quality score, or noise in estimated repeat copy number from individual reads. It will only process a family if all members of the trio have genotypes remaining after filtering. Full documentation of these and other options can be found at https://github.com/gymreklab/STRDenovoTools.

### *SISTR model and method details*

In this study, we also developed SISTR (Selection Inference on Short Tandem Repeats), a novel method to measure negative selection at short tandem repeats. This method is used in the main text **(Fig. 4**) to predict the pathogenicity of specific TR alleles arising from *de novo* mutations. SISTR currently only supports short TRs (STRs) with repeat lengths of 2-4bp, as these repeats are abundant and can be genotyped relatively accurately, and our mutation models are most accurate for these loci[6].

#### Modeling STR mutations

SISTR models STR mutations using a generalized stepwise mutation model (GSM)[7] with two modifications (**Extended Data Fig. 9a**). First, we use a length-dependent mutation rate, given the consistently observed relationship between STR allele length and mutation rate[8-10]. Second, we incorporate a directional bias in mutation sizes, in which short alleles are more likely to expand and long alleles are more likely to contract, consistent with previous observations[9,11]. In our model, this bias pushes alleles back toward a central allele equal to the most frequent population allele length, which for convenience we set to size 0.

The mutation model is characterized by four parameters. The first three: $\mu_0$ (per-generation mutation rate of the central allele), $p$ (mutation step size geometric distribution parameter), and $\beta$ (strength of mutation size directional bias) are the same as in MUTEA[6]. The fourth, $L$, gives the slope of the increase in $\log_{10}$ mutation rate with allele size.

We used our mutation model to build a mutation matrix $M$ that gives the probability of transitioning from allele $g$ to allele $h$ in a single generation for all possible $g$ and $h$. $M$ is an $n\ x\ n$ matrix, where $n$ is the number of possible alleles at the locus, which is set to 25 for all analyses. The central, or "optimal" allele is set as allele 0. The other alleles are assigned nonzero values based on the number of repeats away from the central allele. Let $a_t$ be the value of an allele at generation $t$. The probability of transitioning from allele $a_t = g$ to $a_{t+1} = h$ is given by:

$$P(a_{t+1}|a_t = g) = \begin{cases} 1 - \mu_g, h = g \\ \mu_g f_i P_X(h - g), h > g \\ \mu_g f_d P_X(g - h), h < g \end{cases}$$

Where:

- $\mu_g$ is the mutation rate of allele $g$, computed as $\mu_g = 10^{Lg + \log_{10} \mu_0}$
- $f_i$ and $f_d$ give the probability of a mutation resulting in an increase or decrease in repeat number, respectively. $f_i = \frac{1-\beta pg}{2}$ ; $f_d = 1 - f_i$. For cases where $f_i < 0.01$, we set $f_i = 0.01$ and $f_d = 0.99$. Similarly, for $f_i > 0.99$, we set $f_i = 0.99$ and $f_d = 0.01$ to allow for a minimal probability of expansions or contractions at very large or very small alleles, respectively.
- $P_X(k)$ gives the probability of a mutation resulting in a change in $k$ repeat units and is computed using the probability mass function of a geometric distribution: $P_X(k) = p(1-p)^{k-1};\ k = 1,2,3, \dots$ .

**Modeling selection at STRs**

We modeled selection using a diploid fitness surface (see example in **Extended Data Fig. 9b**). The fitness $(w)$ of each allele $a$ (where here $a$ is the absolute number of repeat units away from the central allele) is given as a function of a selection coefficient $(s)$: $w(a) = 1 - sa$. These per-allele values are then used to calculate the fitness of each genotype to construct the diploid fitness surface. Under an additive model, the fitness of the genotype consisting of alleles $a$ and $b$, $w(a,b)$, is given by half the sum of the fitness of the individual alleles $\frac{w(a)+w(b)}{2}$, with negative genotype fitness values set to 0.

**Forward simulation of allele frequencies**

To model the evolution of STRs, including mutation, negative selection and genetic drift, we simulated allele frequencies forward in time using a previously described algorithm[12] with the addition of an end sampling step based on the number of individuals in the empirical data (see example in **Extended Data Fig. 9c**). The algorithm proceeds as follows:

**Step 0.** Set $t = 0$ and let $\vec{p}_0$ be the $1 \, x \, n$ vector of initial allele frequencies. We set the value to 1 for the central "optimal" allele and 0 for all other alleles. Here $n$ is the number of total possible alleles allowed in the simulation. In all cases, we set $n = 25$.

**Step 1.** Calculate the allele frequencies after one generation of selection and mutation ($\vec{p}_{t+1}$) using the recursion equation $\vec{p}_{t+1} = M^T(\vec{p}_t + \frac{C}{2\bar{w}} \nabla\bar{w})$, where:

- $M^T$ is the transpose of the mutation matrix described above.
- $C$ is the $n \, x \, n$ covariance matrix, where diagonal elements are set to $C[i,i] = p_i(1 - p_i)$, off-diagonal elements are set to $C[i,j] = -p_i p_j$, and $p_i$ is the frequency of allele $i$.
- $\nabla\bar{w}$ is the $n \, x \, 1$ gradient vector of partial derivatives of $\bar{w}$, where $\nabla\bar{w}_i = \frac{\partial\bar{w}}{\partial p_i} = 2w^*(i)$. Here $w^*(i)$ is the marginal fitness of allele $i$ and is computed as $w^*(i) = \sum_j p_j w(i,j)$. Recall from above $w(i,j)$ is the diploid fitness of genotype $i,j$.
- $\bar{w}$ is the mean fitness (the sum of genotypic fitnesses weighed by the corresponding genotype frequency)

$$\bar{w} = \sum_i \sum_j p_i p_j w(i,j) = \sum_i p_i w^*(i)$$

**Step 2.** Recalculate $\vec{p}_{t+1}$ after reproduction and genetic drift. Using the frequencies of each allele in $\vec{p}_{t+1}$ from step 1 as probabilities, use multinomial sampling to draw a sample of size $2N_e$ to obtain a recalculated $\vec{p}_{t+1}$. $N_e$ denotes the effective diploid population size.

This may either be set to a constant value or varied at each time point to reflect realistic demographic histories including population size changes (see a description of demographic models below).

**Step 3.** Repeat Steps 1 and 2 for $g$ generations.

**Step 4.** After $g$ generations, perform an end sampling step. Using the probabilities in $\vec{p}_g$, use multinomial sampling to draw a sample of size $N$ to obtain the final allele frequencies, where $N$ is the number of alleles observed in empirical allele frequencies.

In all cases we run for 50,000 generations, followed by 5,920 generations of population size changes according to the European demographic model described below, for a total of $g = 55{,}920$.

**Estimating selection coefficients using ABC**

We used ABC to obtain a posterior distribution of the selection coefficient $s_i$ for each STR $i$. For each STR, we take the observed allele frequencies, mutation model parameters $(\mu_0, \beta, p, L)$, simulation algorithm parameters $(n, g, N_e, N)$, and parameters $a, b$ describing a gamma distribution prior on $s_i$ (**Extended Data. Fig. 9d**). We then repeat the following steps $z = 10{,}000$ times:

1. Draw a selection coefficient $s$ from the prior $s_i \sim \Gamma(a, b)$.
2. Simulate allele frequencies forward in time in a population using the mutation model and simulation algorithm parameters.
3. Compare simulated to observed allele frequencies using two different summary statistics. The first summary statistic is heterozygosity. The second summary statistic is a $1x5$ vector **v** of allele frequencies, where **v**[0] = sum of frequencies of alleles more than one repeat unit less than the optimal alleles, **v**[1] = frequency of the allele one repeat unit less than the optimal allele, **v**[2] = frequency of allele 0 (central allele), **v**[3] = frequency of the allele 1 repeat unit greater than the optimal allele, and **v**[4] = sum of frequencies of alleles more than one repeat unit greater than the optimal allele. If the simulated and observed summary statistics are

sufficiently close to each other, accept the $s$ value and include it in the posterior distribution. Otherwise, reject the value of $s$ drawn from the prior. For the heterozygosity to be deemed sufficiently close, the difference between the observed and simulated heterozygosity must be less than (observed heterozygosity + 0.005)/3. For the vector **v** of allele frequencies to be considered sufficiently close, the difference between the observed vector $v_o$ and the simulated vector $v_s$ must be less than 0.3.

Values of $s_i$ from all accepted simulations make up the posterior distribution. We report the median of the posterior distribution with a 95% credible interval.

In addition to reporting information on the posterior distribution of $s_i$, we perform a likelihood ratio test (LRT) of whether a model with selection (represented by $s_i = \widehat{s_i}$, where $\widehat{s_i}$ is the ABC median posterior estimate for $s_i$) fits better than a model without selection ($s_i = 0$). Let $L_0 = Likelihood\ (s_i = 0)$ and $L_1 = Likelihood\ (s_i = \widehat{s_i}\ )$. We find the likelihood of $s_i = \widehat{s_i}$ by performing $z = 2{,}000$ simulations given that $s_i$ and determining the fraction of simulations that are sufficiently similar (based on the criteria outlined in step 3 of the ABC procedure above) to the observed allele frequency distribution. Then, the log-likelihood ratio, which is given by $LogLR = -2\ln\frac{L_0}{L_1}$, should asymptotically follow a chi-square distribution with 1 degree of freedom. Here, since the null hypothesis $s = 0$ falls on the boundary of the parameter space, the null consists of a mixture distribution of a point mass at 0, with 50% probability, and a chi-square distribution with 1 degree of freedom, also with 50% probability[13]. We use this mixture distribution to obtain a p-value for each TR based on the log-likelihood ratio.

**SISTR implementation**

To improve computational efficiency, rather than performing a separate set of simulations required for ABC at each locus, we generated one ABC and one LRT lookup table for each of 23 STR classes (an STR's class is determined by its period [e.g. dinucleotide or trinucleotide repeats] and optimal allele [e.g. 6 repeat units]). Each STR class corresponds to a different mutation model with different $p$, $L$, and $\mu_0$values (note: for all

STR classes, $\beta = 0.3$). For dinucleotides, $p = 0.6$, $L = 0.15$, and $\mu_0 = 10^{-5}$ at 6 repeat units; for trinucleotides, $p = 0.9$, $L = 0.33$, and $\mu_0 = 10^{-5.5}$ at 5 repeat units; and for tetranucleotides, $p = 0.9$, $L = 0.45$, and $\mu_0 = 10^{-6}$ at 3 repeat units. These mutation model parameters are based on observed mutations (**Extended Data Fig. 2b**). For each ABC lookup table, we drew $z = 10{,}000$ *s* values from a gamma distribution prior. Then, using the mutation model corresponding to the STR class, we simulated allele frequencies forward in time for each *s* value. Thus, each ABC lookup table contains a list of $s$ values and their corresponding simulated allele frequency distributions.

For each LRT lookup table, for each $s$ value in a list of 45 *s* values (0, 1e-06, 1e-05, 2e-05, 3e-05, 4e-05, 5e-05, 6e-05, 7e-05, 8e-05, 9e-05, 0.0001, 0.0002, 0.0003, 0.0004, 0.0005, 0.0006, 0.0007, 0.0008, 0.0009, 0.001, 0.002, 0.003, 0.004, 0.005, 0.006, 0.007, 0.008, 0.009, 0.01, 0.02, 0.03, 0.04, 0.05, 0.06, 0.07, 0.08, 0.09, 0.1, 0.15, 0.2, 0.4, 0.6, 0.8, 1.0), we simulated allele frequencies forward in time using the $s$ value and the mutation model for the STR class. Simulated results (**Fig. 4a**) are based on LRT lookup tables with 200 simulated allele frequency distributions for each *s* value. Genome-wide results (**Fig. 4b-d**) are based on expanded LRT lookup tables with 2,000 simulated allele frequency distributions per parameter combination.

Based on the period and optimal allele of the allele frequency distribution for which we are estimating *s*, we select the corresponding ABC and LRT lookup tables. Next, we iterate through the ABC lookup table and determine whether to accept each $s$ based on whether the simulated and observed allele frequencies are sufficiently similar to each other based on summary statistics described above. After obtaining a posterior estimate of $s$, we round this *s* value to $s_{round}$ based on the nearest value in 45 $s$ values in the LRT lookup table listed above and set all *s* values less than $10^{-5}$ to 0. Then, from the LRT lookup table, we estimate the likelihood of a particular $s_{round}$ $(L_1 = Likelihood\ (s_i = s_{round}))$ as the percent of simulated allele frequencies for that value that are sufficiently close to the observed allele frequencies. We similarly obtain $L_0 = Likelihood\ (s_i = 0)$ and use these values to compute $LogLR = -2ln\frac{L_0}{L_1}$ which is used to obtain a p-value as described above.

Lookup tables were generated using $n = 25$ alleles, running simulations for $g = 55920$ generations, and end sampling size $N = 6500$. The diploid effective population size for each multinomial sampling step was based on a European demographic model [14,15] described below. The prior from which $s$ was drawn for ABC was a gamma distribution with $a = 0.0881$ and $b = 0.2541$. We found that our results are generally robust to the choice of prior and demography used (**Extended Data Fig. 10c**).

**Demographic models used in SISTR**

SISTR uses a European demographic model based previously published models[14,15]. The population size for each generation in the demographic model is used for each multinomial sampling step in the forward simulation algorithm. The model starts with an ancestral population of size 7,310. Then, after 50,000 generations, the population goes through expansions and contractions over time, including an expansion 5,920 generations ago to a population size of 14,474; an out of Africa bottleneck event 2,040 generations ago to a population size of 1,861; and a European/Asian divergence event 920 generations ago to a population size of 1,032. These events are then followed by 2 periods of recent exponential growth. From 920 to 205 generations ago, the population grew at a rate of 0.00307; from 205 generations ago to present, the growth rate increased to 0.0195.

### Supplementary Note

#### *Enrichment of common variant risk*

We repeated our mutation burden analysis restricting to TRs within 50kb of SNPs previously associated with ASD or related traits through genome-wide association studies (GWAS). We observed no significant mutation excess for TRs near ASD GWAS signals, but observe nominally significant increased mutation burden in ASD probands for mutations near GWAS signals for schizophrenia (SCZ) and educational attainment (EA), which are positively genetically correlated with ASD[16] (**Extended Data Fig. 5a**). The burden is strongest and significant after multiple hypothesis correction for EA (Mann Whitney two-sided p=0.0073) when only considering mutations resulting in common alleles (frequency >0.05; **Extended Data Fig. 5b**), suggesting counter-intuitively that some *de novo* TR mutations may result in ASD risk alleles that are common in the population and may be in linkage disequilibrium with signals identified by SNP-based GWAS. The observation of stronger enrichment for SCZ and EA is consistent with previous analyses of *de novo* point mutations[17] and may be in part due to higher-powered GWAS for those traits compared to ASD.

### *Proband mutations predicted to alter gene expression*

Based on the observed enrichment in fetal brain promoters and previously demonstrated role of TRs in regulating gene expression[18], we hypothesized that *de novo* TR mutations in probands may act in part by altering gene expression during brain development. We examined expression of genes with coding or promoter mutations using the BrainSpan Atlas of the Developing Human Brain[19] resource. We found that genes with TR mutations only observed in ASD probands (proband gene set) show significantly higher prenatal expression compared to genes with only mutations found in unaffected siblings (control gene set) (p=6.3e-15 at 13 post-conceptional weeks [pcw], meta-analysis across 16 brain structures; **Methods; Fig. 3c**). Median expression of the proband gene set is higher across all time points with the strongest effects at prenatal periods (**Extended Data Fig. 6a**) in all brain structures analyzed except cerebellar cortex (CBC) and mediodorsal nucleus of thalamus (MD). We additionally tested whether proband mutations are predicted to alter brain expression of nearby genes based on our previous genome-wide analysis of effects of TR variation on gene expression[18]. We found that predicted effects of proband mutations are significantly stronger than for TRs with only mutations in unaffected siblings in Brain-Caudate (**Extended Data Fig. 7b**; Mann-Whitney two-sided p=0.037), but not for Brain-Cerebellum or the 15 other non-brain tissues analyzed in that study.

We identified specific TR mutations in coding or promoter regions resulting in alleles unobserved in unaffected parents. One example such proband mutation shown in

**Extended Data Fig. 10** is a deletion of 3 copies of CAG in the 5'UTR of *HDAC2*, which results in a previously unobserved allele of 5 copies. In our previous genome-wide analysis of effects of TRs on gene expression[18], we identified a negative association between CAG copy number and expression of *HDAC2*. The *de novo* allele of 5 copies in the proband is thus predicted to increase expression of *HDAC2*, which is highly expressed prenatally in the dorsolateral prefrontal cortex and other brain regions in the BrainSpan database. Notably, a recent study found that *HDAC2* is upregulated in the prefrontal cortex of a Shank3-deficient mouse model of ASD[20] and that down-regulating *HDAC2* rescues social deficits. These results are consistent with the hypothesis that deletion of CAG copies, resulting in increased *HDAC2* expression, could contribute to an ASD phenotype.

### Supplementary Discussion

In addition to replicating known properties of TR mutations, we find significant biases in TR mutation characteristics arising in maternal vs. paternal germlines which provide insights into general biological mechanisms of TR mutation. Strand slippage during DNA replication is widely considered the predominant driver of TR mutations[21]. However, previous studies have reported that other mechanisms, including non-homologous end joining (NHEJ) of DNA double-stranded breaks[22] and recombination-mediated processes such as meiotic gene conversion[23], may also play a role. Intriguingly, we find that mutations derived from maternal germlines are significantly larger and more prone to expansion than those from paternal germlines. Whereas spermatogonia undergo more frequent mitosis events which may lead to a higher rate of slippage events, oocytes lie dormant for decades and can accumulate DNA damage that must be repaired by error-prone processes such as homologous recombination or NHEJ[24]. Further, oocytes have been reported to have crossover frequencies 1.7x of that of spermatocytes[25], providing increased potential for recombination-mediated mutations. Our results are consistent with a stronger influence of slippage resulting in smaller mutations in the paternal germline and of alternative TR mutation processes leading to larger mutations in the maternal germline.
